# Avoidance of MAIT cells is an essential determinant of *Listeria monocytogenes* pathogenesis

**DOI:** 10.1101/2025.10.13.682223

**Authors:** Rafael Rivera-Lugo, Jesse Garcia Castillo, Mariya Lobanovska, Eugene Tang, Andrea Anaya-Sanchez, Scott Espich, Sarah A. Stanley, Michel DuPage, Daniel A. Portnoy

## Abstract

Mucosal-associated invariant T (MAIT) cells are among the most conserved and abundant innate-like T cells in humans that recognize microbial-derived riboflavin precursors and elicit potent antimicrobial responses. The foodborne pathogen *Listeria monocytogenes* is a broad host-range facultative intracellular pathogen that lacks the riboflavin biosynthetic pathway, leading us to hypothesize that this deficiency is pathoadaptive and allows the pathogen to evade MAIT cells. Here, we show that *L. monocytogenes* strains engineered to produce riboflavin (*L. monocytogenes*-*ribDEAHT*) are attenuated in wild-type mice but fully virulent in MAIT cell-deficient mice. Infection with *L. monocytogenes*-*ribDEAHT* prompted rapid and robust MAIT cell expansion in multiple tissues and required the cytolytic effector perforin to eliminate infected cells *in vivo* and *in vitro*. We also assessed the therapeutic potential of *L. monocytogenes*-*ribDEAHT-*stimulated MAIT cells in both infectious disease and cancer mouse models. Therapeutic administration of *L. monocytogenes*-*ribDEAHT* provided protection against *Francisella tularensis* in the lungs and inhibited tumor growth even in the absence of CD8^+^ T cells. These findings reveal the importance of MAIT cell evasion during *L. monocytogenes* infection and highlight the therapeutic potential of engineered *L. monocytogenes* to activate and harness MAIT cells for protection against infectious disease and cancer.

**Significance Statement:** *Listeria monocytogenes* is a bacterial pathogen that grows freely in the environment but can become intracellular following ingestion of contaminated food. Although *L. monocytogenes* can synthesize most metabolites required for growth, it lacks the genes necessary to produce riboflavin (vitamin B2), an essential cofactor across all domains of life. We hypothesized that lacking riboflavin biosynthesis allows *L. monocytogenes* to evade mucosal-associated invariant T cells (MAIT cells), which generate potent antimicrobial responses against riboflavin-producing microbes. By engineering *L. monocytogenes* to produce riboflavin, we show that these strains robustly activate MAIT cells and are highly attenuated in wild-type mice, but not in MAIT cell-deficient mice. Furthermore, MAIT cells activated by engineered *L. monocytogenes* provided therapeutic protection against other riboflavin-producing bacteria and cancer.

## Introduction

MAIT cells are innate-like T cells that primarily recognize a conserved microbial-derived riboflavin precursor presented by monomorphic major histocompatibility complex (MHC) class I-related (MR1) molecules, in contrast to conventional T cells which recognize diverse peptide antigens displayed by polymorphic MHC molecules (1, 2). The MAIT cell semi-invariant T cell receptor (TCR) alpha-chain and MR1 are highly conserved among nearly all mammals, suggesting strong evolutionary pressure for their maintenance (3). Upon activation, MAIT cells produce potent antimicrobial responses, including pro-inflammatory cytokines and/or cytotoxic effectors targeting infected host cells (4, 5). MAIT cells also contribute to tissue repair and protection against a wide range of pathogens, including viruses, riboflavin-producing bacteria, and fungi. In humans, MAIT cell deficiency has been associated with susceptibility to bacterial and viral infections, immune disorders and some cancers (6, 7). The evolutionary conservation and robust effector functions of MAIT cells highlight their therapeutic potential to combat difficult-to-treat bacterial infections and cancer.

*L. monocytogenes* is a facultative intracellular bacterial pathogen that is ubiquitous in the environment and infects many mammals including humans where it grows in the host cell cytosol (8). *L. monocytogenes* can synthesize most of its own required metabolites (9), but unlike most bacteria, it lacks the capacity for *de novo* riboflavin synthesis (10, 11) and instead imports the essential riboflavin-derived flavin mononucleotide (FMN) and flavin adenine dinucleotide (FAD) cofactors from the host cell cytosol using its RibU transporter (12) (**Fig. 1A**). These findings raise a fundamental question: if flavins are essential for *L. monocytogenes* metabolism, survival, and pathogenesis (12–14), does the inability to synthesize riboflavin provide a selective advantage? We hypothesized that the absence of the riboflavin biosynthetic pathway allows *L. monocytogenes* to avoid detection by MAIT cells. To test this hypothesis, we engineered *L. monocytogenes* to produce riboflavin (*L. monocytogenes*-*ribDEAHT*) and investigated the role of MAIT cells during infection. We found that MAIT cells robustly proliferated in infected tissues and restricted the growth of *L. monocytogenes*-*ribDEAHT,* suggesting that scavenging of flavin cofactors instead of riboflavin biosynthesis provides *L. monocytogenes* with a selective advantage in animals that have MAIT cells.

**Figure 1.**
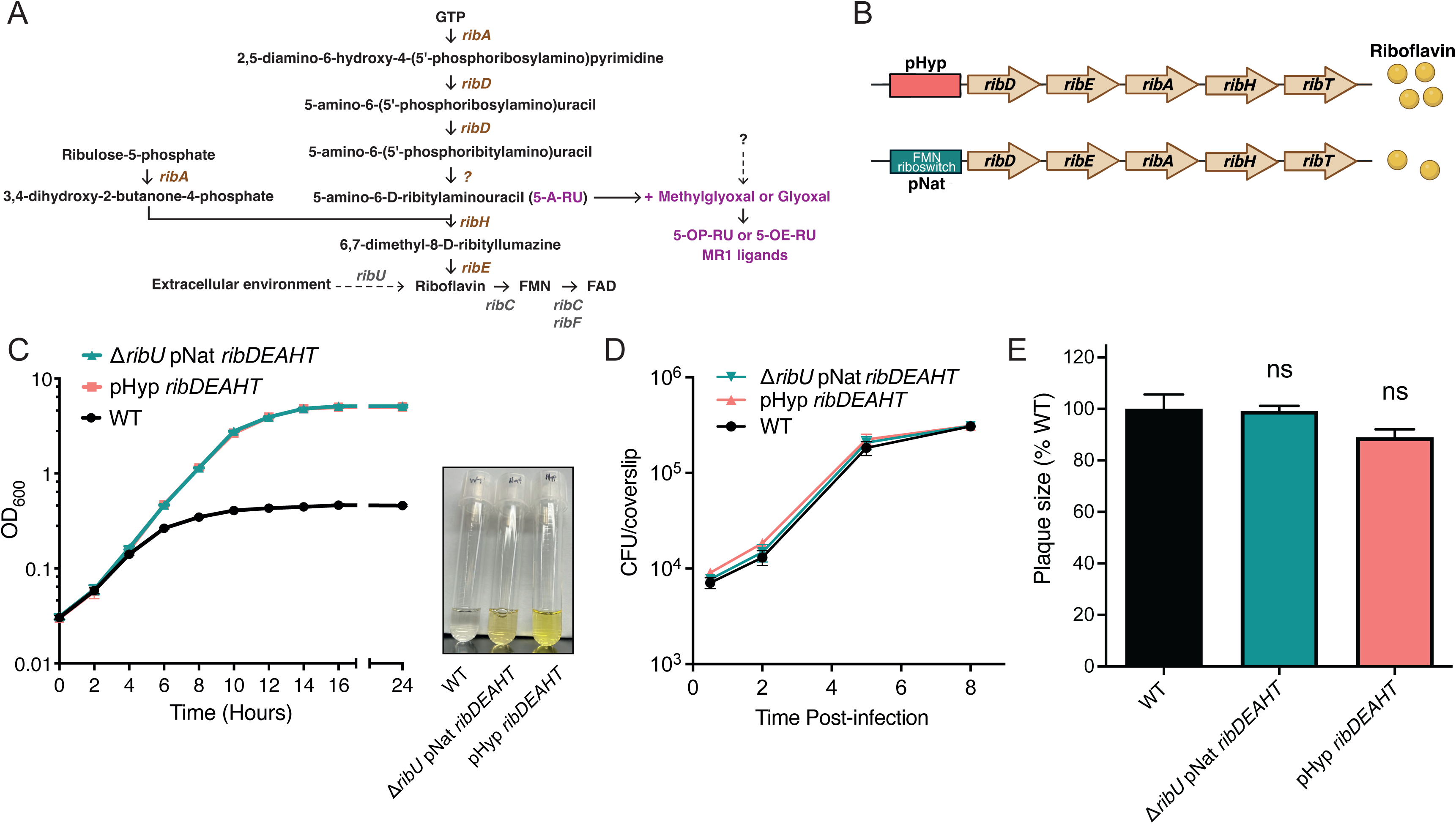
*L. monocytogenes* expressing *ribDEAHT* produce riboflavin and have no detectable virulence defects *in vitro*. (A) Schematic diagram of riboflavin biosynthesis genes (83, 84). *L. monocytogenes* specific genes are shown in grey, *B. subtilis* genes are shown in brown. (B) Schematic of *B. subtilis ribDEAHT* operon used for construction of the riboflavin-producing *L. monocytogenes* strains. *ribDEAHT* under constitutive promoter (pHyp) and under native *B. subtilis* promoter containing an FMN riboswitch (pNat). (C) Growth of indicated *L. monocytogenes* strains in chemically defined media lacking flavins at 37°C with agitation. Cell density was measured by optical density (OD_600_) at the indicated time points. Means and SD from three experiments are shown and the image was taken of the supernatant of the indicated strains at 24 hour of growth. The change in color from colorless to bright yellow (natural color of flavins) is indicative of riboflavin production. (D) Intracellular growth curves of *L. monocytogenes* strains in murine BMMs infected with indicated strains at a multiplicity of infection (MOI) of 0.25, CFUs were enumerated at indicated time points. Gentamicin (50 μg/mL) was added at 1 hour post-infection. Data show means and SEM of three experiments. (E) Plaque formation of *L. monocytogenes* strains in rat L2 fibroblast monolayers. Mean and SD plaque size of each strain from three independent experiments is shown as a percentage relative to WT plaque size. Statistical analysis was performed by one-way ANOVA and Dunnett’s post-test using WT as control. ns, not significant, P > 0.05.

Despite its pathogenic potential, highly attenuated *L. monocytogenes* strains have been developed as vaccine vectors for cancer immunotherapy, stemming from their capacity to robustly induce CD8^+^ T cells (15). In this study, we evaluated the therapeutic potential of recombinant, attenuated *L. monocytogenes*-*ribDEAHT* that induce MAIT cells. Remarkably, riboflavin-producing strains of *L. monocytogenes* expanded and activated MAIT cells that infiltrated the lungs and tumors and protected against *Francisella tularensis* infection, and suppressed tumor growth in the absence of CD8^+^ T cells.

## Results

### *L. monocytogenes* engineered to produce riboflavin have no detectable virulence defects in vitro

We hypothesized that by lacking the riboflavin biosynthetic pathway (**Fig. 1A**), *L. monocytogenes* evades MAIT cell activation and responses. To test this hypothesis, we engineered *L. monocytogenes* strains capable of synthesizing riboflavin *de novo* by introducing a five-gene riboflavin biosynthetic operon (*ribDEAHT)* from the related Gram-positive bacterium *Bacillus subtilis* (12, 13, 16). *ribDEAHT* was introduced in wild-type (WT) *L. monocytogenes* either under a constitutive promoter (pHyp *ribDEAHT*) or under its native *B. subtilis* promoter (pNat *ribDEAHT*), which is negatively regulated by intracellular flavin levels (16) (**Fig. 1B**). To maximize riboflavin synthesis in *L. monocytogenes* pNat *ribDEAHT*, the gene encoding the flavin importer (*ribU*) was deleted, resulting in Δ*ribU* pNat *ribDEAHT*. Both *L. monocytogenes* pHyp *ribDEAHT* and Δ*ribU* pNat *ribDEAHT* displayed robust growth in chemically defined synthetic media in the absence of exogenous flavins suggesting that both strains produced riboflavin (**Fig. 1C**). Riboflavin synthesis from *ribDEAHT* expression changed the color of the media from translucent to bright yellow (**Fig. 1C**), the natural color of flavins (10). The color intensity between the strains varied, with the pHyp *ribDEAHT* producing a higher color intensity. These results implied that although both strains produced riboflavin *de novo*, pHyp *ribDEAHT* resulted in more riboflavin production than Δ*ribU* pNat *ribDEAHT*.

To determine if riboflavin-producing *L. monocytogenes* exhibited MAIT cell-independent virulence defects we first characterized the growth of the strains in *in vitro* infection models in the absence of MAIT cells. *L. monocytogenes*-*ribDEAHT* strains grew intracellularly in bone marrow-derived macrophages (BMMs) and formed plaque sizes in L2 fibroblasts similar to WT bacteria (**Fig. 1D, 1E**). Collectively these data clearly showed that riboflavin-producing *L. monocytogenes* infected, escaped from phagosomes, grew intracellularly, and spread cell-to-cell similar to WT *L. monocytogenes*. These findings indicate that riboflavin biosynthesis does not impair bacterial fitness *in vitro*, allowing us to assess its impact on virulence *in vivo*. As shown below, the riboflavin-producing *L. monocytogenes* strains are attenuated in WT mice but fully virulent in mice lacking MAIT cells, revealing that this phenotype is dependent on MAIT cells.

### MAIT cells restrict riboflavin-producing *L. monocytogenes* strains during murine infection

Next, we evaluated bacterial growth dynamics in a standard murine model of infection. C57BL/6J mice were infected intravenously (i.v.) with 10^3^ colony-forming units (CFUs) of WT or riboflavin-producing *L. monocytogenes* strains and bacterial growth was monitored over the course of 7 days (**Fig. 2A**). Riboflavin-producing strains displayed minor or no virulence defect early during infection (2 days) but were significantly attenuated compared to WT throughout the subsequent course of infection. Both pHyp *ribDEAHT* and Δ*ribU* pNat *ribDEAHT* strains exhibited a similar pattern of growth early during infection in spleens and livers. At 7 days post-infection the CFUs of WT and *L. monocytogenes*-*ribDEAHT* strains dropped in spleens and livers (**Fig. 2A**). However, the virulence defect of the *ribDEAHT*-containing strains was more pronounced in spleens with no CFUs of riboflavin-producing strains detected in the spleens at 7 days post-infection. In livers between 4 and 7 days of infection Δ*ribU* pNat *ribDEAHT* remained attenuated with no changes in CFUs whereas pHyp *ribDEAHT* CFUs continued to drop, suggesting differences in the growth kinetics between the two strains. The decreased infectivity of the *L. monocytogenes*-*ribDEAHT* strains observed *in vivo*, but not *in vitro* (**Fig. 1D-E**), suggested that riboflavin synthesis was detrimental to *L. monocytogenes* virulence. To test whether riboflavin-producing strains resulted in MAIT cell expansion, we examined the percentage of MAIT cells identified as CD3^+^CD4^−^CD8^−^(DN), MR1:5-OP-RU tetramer^+^ T cells in both spleens and livers at 4 days post-infection, since the largest difference in CFUs between the strains was observed at 4 days (**Fig. 2A**). In naïve controls or in WT *L. monocytogenes* infected organs, MAIT cells constituted 0.25-0.45% of DN αβ T cells. The frequency of MAIT cells in mice was approximately 5-fold higher in spleens infected with *L. monocytogenes*-*ribDEAHT* strains and 3-fold higher in livers infected with Δ*ribU* pNat *ribDEAHT* (**Fig. 2B**). These data supported the conclusion that MAIT cells responded specifically to riboflavin-producing *L. monocytogenes* and accumulated in infected tissues.

**Figure 2.**
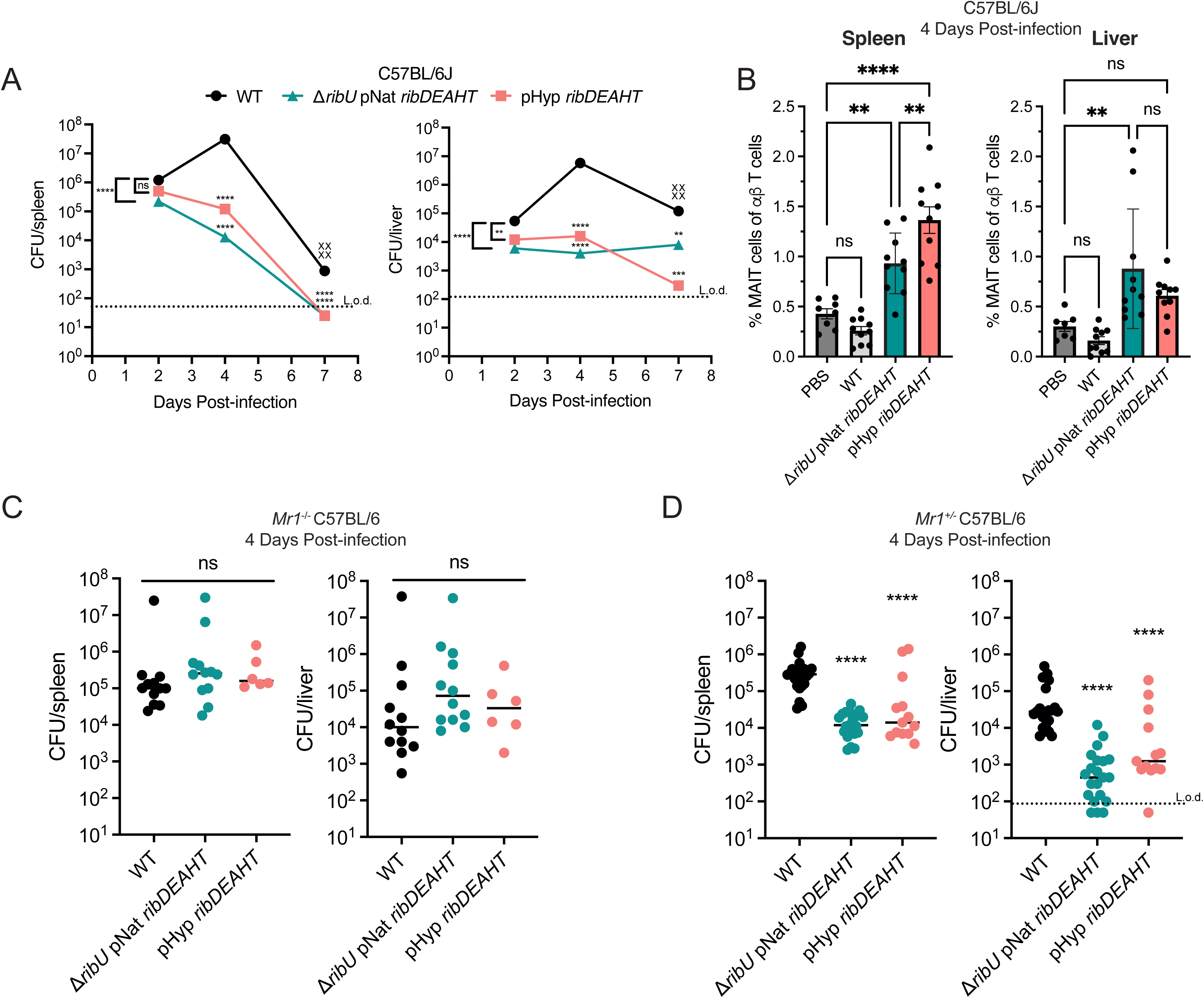
*L. monocytogenes* strains producing riboflavin are attenuated *in vivo*. (A) Bacterial burdens in spleens (left) and livers (right) of C57BL/6J mice at 2, 4, and 7 days post-infection with 1×10^3^ CFUs of indicated *L. monocytogenes* strains. Data show the median from three independent experiments, *n*=15 mice per time point. “X” indicates mice that succumbed to infection. Statistical significance of log-transformed CFUs values was determined by one-way ANOVA and Dunnett’s post-test using WT as the control for each individual time point. (B) Frequency of MAIT cells in the spleens and livers of C57BL/6J mice at 4 days post-infection with 1×10^3^ CFUs of WT (*n*=10), Δ*ribU* pNat *ribDEAHT* (*n*=10), pHyp *ribDEAHT* (*n*=10), or PBS-infected (*n*=8) analyzed using flow cytometry. Means and SEM of two experiments. One-way ANOVA and Dunnett’s post-test were performed using naïve mice (PBS) as the control. (C and D) Bacterial burdens in spleens and livers 4 days post-infection of (C) *Mr1*^−/−^ and (D) *Mr1*^+/−^heterozygous C57BL/6 littermate controls infected with 1×10^3^ CFUs of indicated *L. monocytogenes* strains. In (C), data show the median of least two independent experiments: WT (*n*=12), Δ*ribU* pNat *ribDEAHT* (*n*=12), and pHyp *ribDEAHT* (*n*=6). In (D), data represent the medians of three experiments: WT (*n*=22), Δ*ribU* pNat *ribDEAHT* (*n*=20), and pHyp *ribDEAHT* (*n*=13). Dashed lines indicate the limit of detection (L.o.d). One-way ANOVA and Dunnett’s post-test were performed using WT as the control. ****P < 0.0001; ***P < 0.001; **P < 0.01; ns, not significant, P > 0.05.

To test if MAIT cells contributed to the reduction of the bacterial burden of the *L. monocytogenes*-*ribDEAHT* strains, we infected MR1 knockout (*Mr1^−/−^*) mice, which lack MAIT cells, and MR1 heterozygous (*Mr1^+/−^*) littermate controls (17, 18). No differences in CFUs were observed between riboflavin-producing strains and WT *L. monocytogenes* in the spleens or livers of *Mr1^−/−^* mice (**Fig. 2C**). In contrast, in *Mr1^+/−^*mice, *L. monocytogenes*-*ribDEAHT* strains exhibited 1-log and 2-log virulence defects in spleens and livers, respectively (**Fig. 2D**). These results strongly suggested that MAIT cells were solely responsible for decreasing the bacterial burden of the *L. monocytogenes*-*ribDEAHT* strains *in vivo*. Comparison between pHyp *ribDEAHT* and Δ*ribU* pNat *ribDEAHT L. monocytogenes* infections revealed that MAIT cell expansion was marginally different between the two strains in spleens and comparable in livers (**Fig. 2B**), suggesting that *in vivo* both strains may produce similar amounts of ligand for MAIT cell activation. Considering that both strains activated MAIT cells similarly, we decided to focus our subsequent analyses on *L. monocytogenes* strains containing pHyp *ribDEAHT* which produces more riboflavin *in vitro* (**Fig. 1C)**.

### Attenuated *L. monocytogenes*-*ribDEAHT* strains generate robust expansion and persistence of MAIT cells in infected tissues

Based on the observation that MAIT cell expansion correlated with ligand availability (19), we hypothesized that the observed expansion of MAIT cells (**Fig. 2B**) could be further enhanced by increasing the bacterial infectious dose, thereby providing more *L. monocytogenes*-*ribDEAHT*-derived MAIT cell ligand. However, since higher *L. monocytogenes* bacterial loads are lethal, we introduced pHyp *ribDEAHT* into a well-characterized highly attenuated *L. monocytogenes* strain that cannot spread from cell-to-cell (Δ*actA*) but retains potent CD8^+^ T cell-dependent immunogenic potential (20). C57BL/6J mice were infected with 10^7^ CFUs of Δ*actA* or Δ*actA* pHyp *ribDEAHT* and CFUs were assessed 4 days post-infection. The Δ*actA* pHyp *ribDEAHT* strain was at least 1,000-fold less virulent than Δ*actA*, with over 5-log and 3-log virulence attenuation in spleens and livers respectively (**Fig. 3A**). Strikingly, the frequency of MAIT cells in Δ*actA* pHyp *ribDEAHT* infected mice exceeded 15% and 25% of total αβ T cells in spleens and livers, respectively, compared to less than 1% of MAIT cells present following phosphate-buffered saline (PBS) or Δ*actA* infection (**Fig. 3B**).

**Figure 3.**
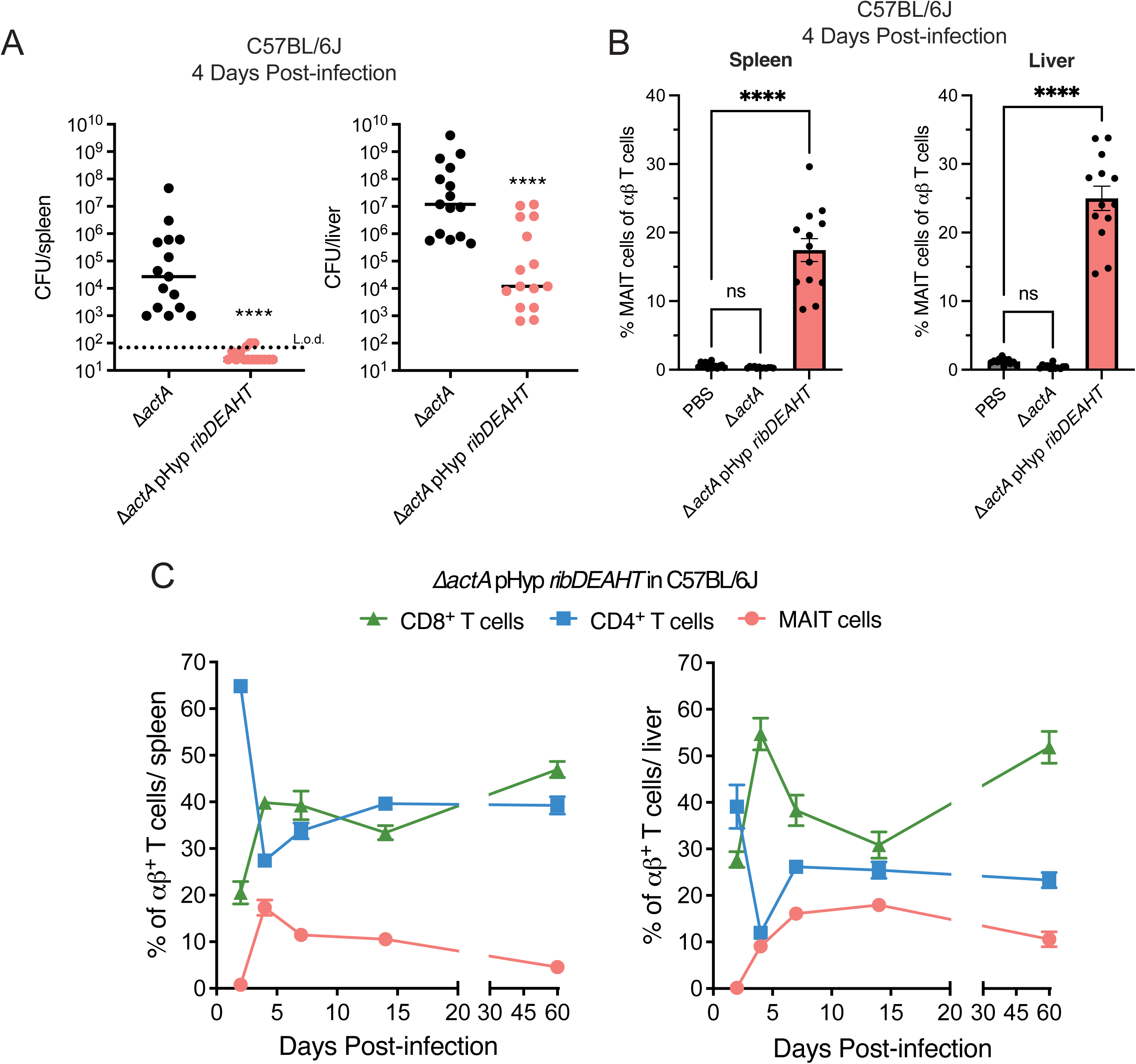
Infection with attenuated *L. monocytogenes*-*ribDEAHT* strains leads to robust expansion and sustained accumulation of MAIT cells in infected tissues. (A) Bacterial burdens 4 days post-infection in spleens (left) and livers (right) of C57BL/6J mice infected with 1×10^7^ CFUs of indicated *L. monocytogenes* strains. Data present the median CFUs, *n*=15 mice per group from three independent experiments. Dashed line is limit of detection (L.o.d.). Statistical significance of log-transformed CFUs values was determined by one-way ANOVA and Dunnett’s post-test using WT as the control. (B) Frequency of MAIT cells in spleens and livers of C57BL/6J mice at 4 days post-infection with 1×10^7^ CFUs of Δ*actA*, Δ*actA* pHyp *ribDEAHT*, or PBS-infected, determined by flow cytometry. Data represent the means and SEM from three independent experiments: naive (*n*=14), Δ*actA* (*n*=10), and Δ*actA* pHyp *ribDEAHT* (*n*=13). Statistical significance was determined using one-way ANOVA and Dunnett’s post-test using naïve mice as the control. (C) αβ T cell kinetics in spleens (left) and livers (right) of C57BL/6J mice showing the frequencies of MAIT, CD4^+^, and CD8^+^ T cells at 2, 4, 7, 14, and 60 days post-infection with 1×10^7^ CFUs of Δ*actA* pHyp *ribDEAHT.* Data represents two independent experiments with 10 mice per each time point. ****P < 0.0001; ns, not significant (P > 0.05).

MAIT cells can persist in tissues long after infection resulting in an innate memory-like phenotype (21–23). To determine the kinetics of MAIT cell accumulation, we monitored the frequency of MAIT cells, CD4^+^ T cells, and CD8^+^ T cells following infection with 10^7^ CFUs of Δ*actA* or Δ*actA* pHyp *ribDEAHT* over the course of 60 days (**Fig. 3C, Fig. S1**). At 2 days post Δ*actA* pHyp *ribDEAHT* infection, MAIT cells comprised less than 1% of all αβ T cells in spleens and livers (**Fig. 3C**) but rapidly expanded so that MAIT cell frequencies peaked at approximately 20% on day 4 post-infection in the spleens, and on day 7 in the livers and persisted at elevated levels for 60 days (**Fig. 3C**). At 60 days post-infection, MAIT cells comprised an average of 5% and 10% of all αβ T cells in the spleens and livers, respectively (**Fig. 3C**). CD4^+^ and CD8^+^ T cell frequencies were comparable between Δ*actA* and Δ*actA* pHyp *ribDEAHT* suggesting that the expansion of MAIT cells did not affect relative frequencies of conventional T cells during *L. monocytogenes* infection, although we did not measure *L. monocytogenes-*specific T cells. MAIT cells were below 1% in mice following Δ*actA* infection (**Fig. S1**). These data demonstrated that MAIT cells responded strongly to attenuated riboflavin-producing *L. monocytogenes* strains by accumulating and persisting in infected organs.

### MAIT cells require perforin to restrict riboflavin-producing *L. monocytogenes*

MAIT cells use two major effector functions to control pathogens: cytokine production, which activates bystander cells, and direct killing of infected cells *via* cytolytic effectors granzyme B and perforin (24, 25). We hypothesized that MAIT cells restricted riboflavin-producing *L. monocytogenes* (**Fig. 2**) by killing infected host cells, thereby eliminating the bacterial replicative niche similar to the mechanism of immunity conferred by CD8^+^ T cells (26). To determine whether perforin plays a role in MAIT cell-dependent restriction of riboflavin-producing *L. monocytogenes*, we infected mice lacking perforin (*Prf1^−/−^*) (27) with 10^3^ CFUs of WT or pHyp *ribDEAHT* and measured bacterial burden 4 days post-infection. *L. monocytogenes*-*ribDEAHT* was equally virulent to WT in *Prf1^−/−^* mice, indicating that perforin was required to control *L. monocytogenes*-*ribDEAHT* growth *in vivo* (**Fig. 4A**). To confirm that MAIT cell expansion still occurred in infected tissues of *Prf1^−/−^*mice in response to riboflavin-producing *L. monocytogenes*, we infected *Prf1^−/−^*mice with 10^7^ CFUs Δ*actA L. monocytogenes*-*ribDEAHT* and determined the frequencies of MAIT cells at 4 days post-infection. MAIT cells constituted approximately 13% and 20% of all αβ T cells in the spleens and livers, respectively (**Fig. 4B**), which was similar to the frequencies observed in infected C57BL/6J mice (**Fig. 3B**). Together, these results suggested that *Prf1^−/−^* MAIT cells expanded upon infection with riboflavin-producing *L. monocytogenes* but were unable to restrict the engineered strains in the absence of perforin.

**Figure 4.**
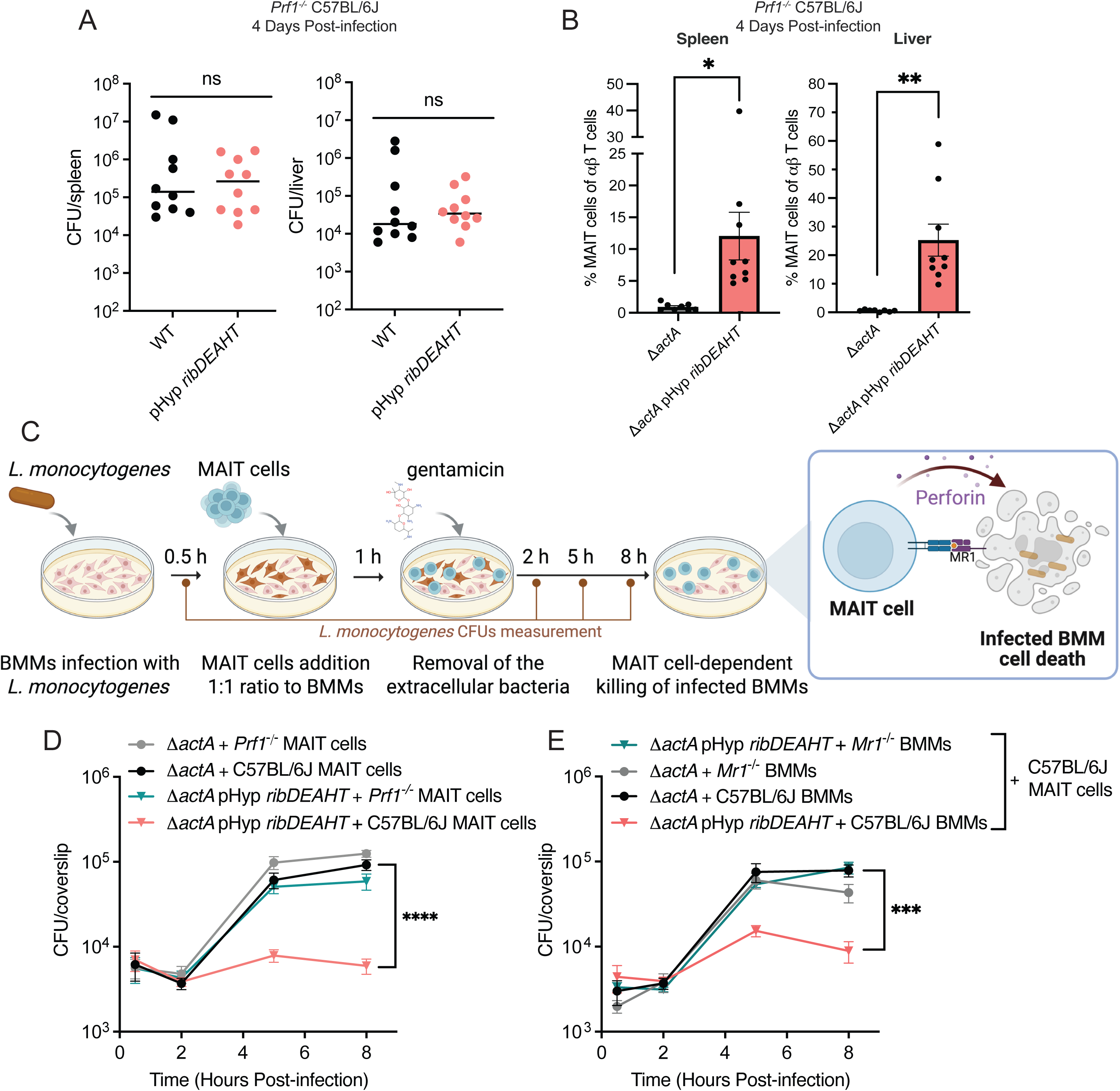
Perforin-deficient MAIT cells do not restrict growth of riboflavin-producing *L. monocytogenes*. (A) Bacterial burdens in spleens (left) and livers (right) of perforin knockout *Prf1*^−/−^ C57BL/6J mice infected with 1×10^3^ CFUs of *L. monocytogenes* strains. Data is from two independent experiments: WT (*n*=10) and pHyp *ribDEAHT* (*n*=10). (B) Frequency of MAIT cells in the spleens and livers of *Prf1*^−/−^ C57BL/6J mice infected with 1×10^7^ CFUs of Δ*actA* or Δ*actA* pHyp *ribDEAHT*. Means and SEM of two independent experiments are shown: Δ*actA* (*n*=8) and Δ*actA* pHyp *ribDEAHT* (*n*=9). One-way ANOVA and Welch’s t-test was performed using WT (A) or Δ*actA* (B) mice as the control. (C) Schematic diagram depicting *in vitro* co-culture system developed to study MAIT cell-dependent cytotoxicity. BMMs infected with *L. monocytogenes* were incubated with *in vitro* expanded MAIT cells and *L. monocytogenes* CFUs were measured during an 8-hour infection. MAIT cell-dependent cytotoxicity of infected BMMs was monitored by measuring the drop of *L. monocytogenes* CFUs. (D and E) Intracellular growth curves of *L. monocytogenes* strains in (D) BMMs co-incubated with MAIT cells from C57BL/6J or *Prf1*^−/−^ mice and in (E) BMMs isolated from C57BL/6J or *Mr1*^−/−^ mice co-incubated with MAIT cells from C57BL/6J. Extracellular bacteria were removed by addition of gentamicin (50 μg/mL) at 1 h post-infection. CFUs were enumerated at the indicated times. Data represent the means and SEM of two (D) and three (E) independent experiments. One-way ANOVA and Dunnett’s post-test were performed using Δ*actA* as control. ****P < 0.0001, ***P < 0.001; **P < 0.01; *P<0.1; ns, not significant (P > 0.05).

There are several populations of perforin-expressing cells including MAIT cells, natural killer (NK) cells and CD8^+^ T cell (24, 28, 29), which may play a role in restricting *L. monocytogenes*-*ribDEAHT* growth *in vivo*. To determine whether MAIT cells are directly responsible for perforin-dependent restriction of riboflavin-producing *L. monocytogenes* strains, we developed an *in vitro* co-culture system based on a previously described method used for studying CD8^+^ T cell cytotoxicity of *L. monocytogenes*-infected macrophages (30). BMMs were infected with *L. monocytogenes* and 30 minutes post-infection were overlaid with *in vitro* expanded MAIT cells derived from spleens from either C57BL/6J or *Prf1^−/−^* mice infected with Δ*actA* pHyp *ribDEAHT*. Intracellular bacterial CFUs were then monitored over the course of 8 hours (**Fig. 4C**). The presence of C57BL/6J MAIT cells had no significant effect on Δ*actA L. monocytogenes* but resulted in a 10-fold drop in Δ*actA* pHyp *ribDEAHT* CFUs between 2- and 8-hours post-infection (**Fig. 4D**). In contrast, *Prf1^−/−^* MAIT cells failed to suppress the growth of Δ*actA L. monocytogenes*-*ribDEAHT* or Δ*actA* strains (**Fig. 4D)**, confirming that MAIT cells required perforin to restrict *L. monocytogenes*-*ribDEAHT* in the BMM growth niche and implying that the mechanism of restriction was perforin-mediated cytotoxicity. Consistent with the direct role of MAIT cells in restricting riboflavin-producing bacteria, a 10-fold reduction in bacterial CFUs was observed when MAIT cells were added to C57BL/6J but not *Mr1*^−/−^ BMMs infected with Δ*actA* pHyp *ribDEAHT* (**Fig. 4E)**. These results suggest that MAIT cell-mediated killing of BMMs infected with riboflavin-producing bacteria was dependent on both perforin and MR1 expression.

### *L. monocytogenes*-*ribDEAHT* stimulated-MAIT cells provide protection against *F. tularensis* infection and restrict tumor growth independent of CD8^+^ T cells

We next considered whether *L. monocytogenes*-*ribDEAHT*-induced expansion, long-term persistence and cytotoxic potential of MAIT cells could be harnessed to stimulate antibacterial and antitumor immunity. First, we examined the vaccination potential of riboflavin-producing *L. monocytogenes* against *F. tularensis* LVS, as *F. tularensis* has been shown to be restricted by MAIT cells in the lungs (31, 32). To determine whether *L. monocytogenes*-*ribDEAHT* infection leads to MAIT cell expansion in the lungs, mice were infected with either Δ*actA or* Δ*actA* pHyp *ribDEAHT* and MAIT cell frequencies were measured in the lungs at 4 days post-infection. MAIT cell frequencies in the lung increased to about 15% of all αβ T cells (**Fig. 5A**), which was similar to the MAIT cell frequencies observed in other tissues (**Fig. 3B**). To assess if *L. monocytogenes*-*ribDEAHT* stimulated-MAIT cells protected against *F. tularensis* LVS lung infection, mice were first infected with 10^6^ CFUs of Δ*actA or* Δ*actA* pHyp *ribDEAHT*. After *L. monocytogenes* clearance on day 14, mice were challenged with 10^4^ CFUs of *F. tularensis* LVS subcutaneously and lungs were harvested on day 19 for *F. tularensis* LVS CFUs enumeration (**Fig. 5B**). Mice vaccinated with Δ*actA* pHyp *ribDEAHT* displayed a more robust control of *F. tularensis* LVS compared to PBS or Δ*actA* vaccinated groups (**Fig. 5C**). Vaccination with Δ*actA* alone conferred 1-log of protection against *F. tularensis* LVS lung infection, while protection in mice vaccinated with the Δ*actA* riboflavin-producing strain was more than 100-fold (**Fig. 5C**). Interestingly, Δ*actA* pHyp *ribDEAHT* vaccination of perforin-deficient mice provided protection against *F. tularensis* comparable to that observed in WT mice (**Fig. S2**), suggesting that the cytotoxic effector functions are not required for the protection response induced by vaccination with Δ*actA* pHyp *ribDEAHT.* Together, these results suggested that vaccination with attenuated *L. monocytogenes*-*ribDEAHT* leads to potent and protective MAIT cell responses against riboflavin-producing pathogens like *F. tularensis*.

**Figure 5.**
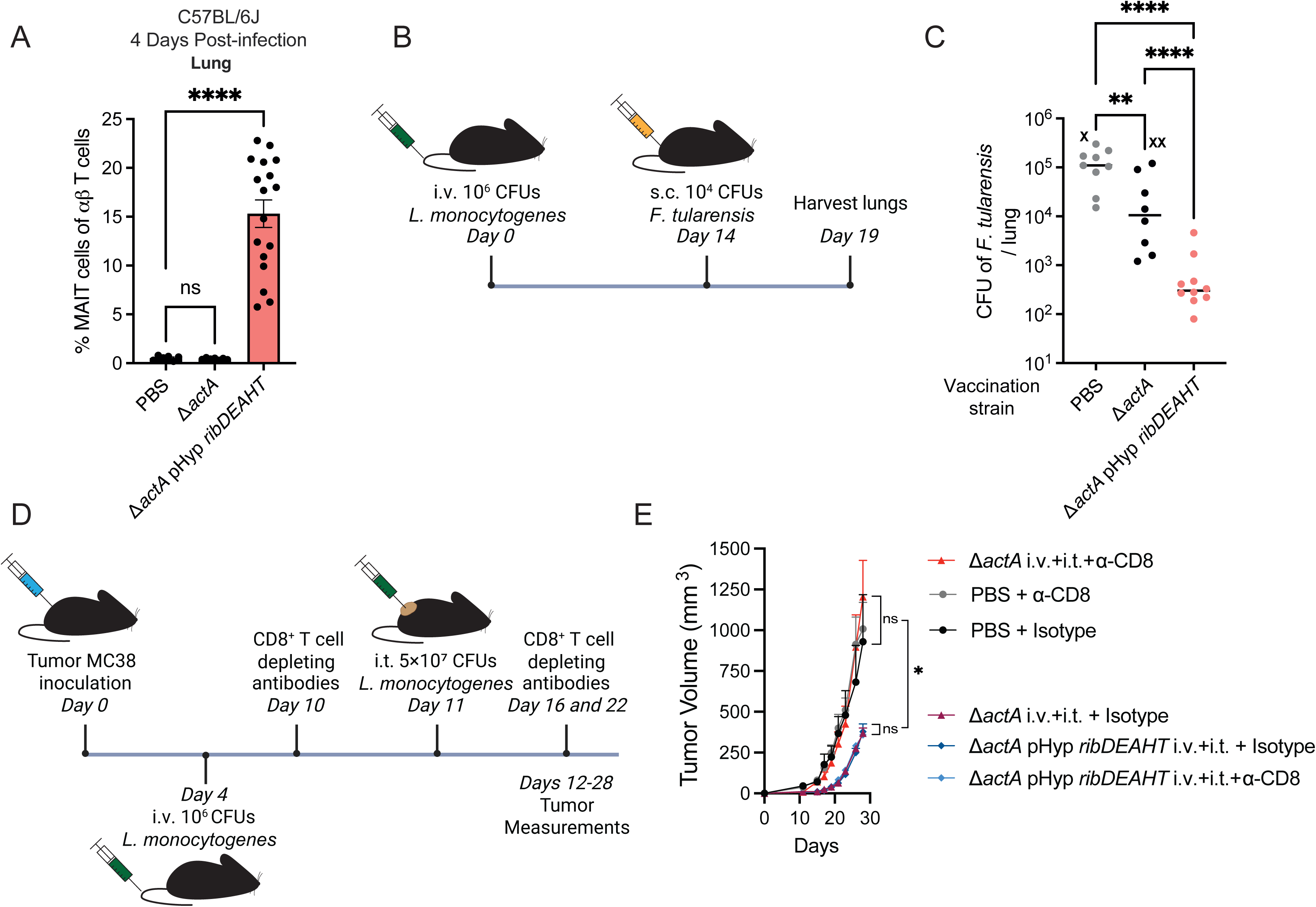
Riboflavin-producing *L. monocytogenes* provides protection against *F. tularensis* infection and inhibit tumor growth. (A) Frequency of MAIT cells in lungs of C57BL/6J mice infected with 1×10^7^ CFUs *L. monocytogenes*. Means and SEM are from two experiments: PBS (*n*=6), Δ*actA* (*n*=7), and Δ*actA* pHyp *ribDEAHT* (*n*=17). One-way ANOVA and Dunnett’s post-test were performed with naïve (PBS) mice as control. (B) Schematic of the vaccination protocol: mice were infected intravenously (i.v.) with 1×10^6^ CFUs *L. monocytogenes* and on day 14 mice were infected with 1×10^4^ CFUs *F. tularensis* LVS subcutaneously (s.c.) and lungs were harvested on day 19. (C) *F. tularensis* LVS lung burdens on day 19 in C57BL/6J mice vaccinated with the indicated *L. monocytogenes* strains. Aggregated data from two experiments: PBS (*n*=9), Δ*actA* (*n*=10), and Δ*actA* pHyp *ribDEAHT* (*n*=10). “x” indicates mice that succumbed to infection. One-way ANOVA and Tukey’s multiple comparisons test was performed. (D) Schematic of tumor protocol: MC38 tumors were implanted in C57BL/6J mice and on day 4 mice were infected i.v. with 1×10^6^ CFUs *L. monocytogenes*. Mice were then infected intratumorally (i.t.) on day 11 and tumors were measured from day 12-28. Mice were treated with either CD8 depleting antibody (anti-CD8b.2) or isotype control on days 10, 16 and 22. (Adapted from (33)) (E) Tumor volume measurements in i.v.+ i.t. treated mice as described in (D). Means and SEM of *n*=5 mice per group, (*n*=3) PBS + isotype control group. Two-way ANOVA and Šídák’s multiple comparisons test were performed using last time point of each group. The representative experiment of three independent biological experiments is shown. ****P < 0.0001; **P < 0.01; *P<0.1; ns, not significant (P > 0.05).

Recombinant attenuated *L. monocytogenes* strains have been used as therapeutic cancer vaccines in preclinical and clinical trials (15). *L. monocytogenes* vaccine strains trigger potent cytotoxic CD8^+^ T cells that infiltrate tumors and mediate remodeling of the tumor microenvironment (33, 34). To investigate whether immunization with *L. monocytogenes*-*ribDEAHT* can lead to MAIT cell expansion in tumors and affect tumor growth, we used a recently described mouse MC38 tumor model that relies on anti-*L. monocytogenes* CD8^+^ T cells for tumor restriction (33) (**Fig. 5D**). Following tumor implantation with MC38 tumor cells, mice were immunized i.v. with *L. monocytogenes* on day 4 and then intratumorally (i.t.) into palpable tumors on day 11. At day 5 following i.v. + i.t. treatment, MAIT cell frequencies increased significantly in tumors infected with Δ*actA* pHyp *ribDEAHT* compared to PBS or Δ*actA* control groups (**Fig. S3**). Next, we assessed whether i.v. + i.t. treatment with Δ*actA* pHyp *ribDEAHT* affected tumor growth. Since the i.v. + i.t. regimen with Δ*actA* requires CD8^+^ T cells to inhibit tumor growth (33), mice infected with Δ*actA* pHyp *ribDEAHT* were treated with CD8-depleting antibodies (α-CD8b.2) or isotype control during i.v. + i.t. treatment (**Fig. 5D**) to determine the contribution of MAIT cells and CD8^+^ T cells to tumor-specific immunity. Remarkably, the tumor-restricting properties of Δ*actA* pHyp *ribDEAHT* were not dependent on CD8^+^ T cells as both CD8 antibody-treated and isotype control groups showed comparable reduced tumor size during i.v. + i.t. treatment (**Fig. 5E**). CD8^+^ T cell-independent tumor restriction was specific to *L. monocytogenes* expressing *ribDEAHT* as tumor growth restriction in Δ*actA* treated mice was abolished when CD8^+^ T cells were depleted (**Fig. 5E**). Together, these results showed that treatment with Δ*actA* pHyp *ribDEAHT* increased tumor-infiltrating MAIT cells and elicited a potent tumor restricting response that does not rely on CD8^+^ T cells.

## Discussion

*L. monocytogenes* synthesizes most of its own metabolites but has an unusual requirement for riboflavin. The results of this study showed that riboflavin dependency is an essential determinant of *L. monocytogenes* pathogenesis, as *L. monocytogenes* strains engineered to produce riboflavin were over 100-fold less virulent in C57BL/6J mice but fully virulent in mice lacking MAIT cells or perforin (**Fig. 2A**, **Fig. 2C**, **Fig. 4A**). Introducing the riboflavin operon into Δ*actA L. monocytogenes,* a well-established vector used in dozens of clinical trials (15), led to robust MAIT cell expansion in the spleens, livers and lungs (**Fig. 3B**, **Fig. 5A**) and provided considerable protection against *F. tularensis* (**Fig. 5C**). Most strikingly, Δ*actA L. monocytogenes-ribDEAHT* triggered expansion of MAIT cells in tumors and prevented tumor growth (**Fig. S3, Fig. 5E**).

Since most bacteria and fungi synthesize riboflavin and mammals do not, it is not at all surprising that an intermediate of riboflavin biosynthesis is a MAIT cell ligand. While some pathogens modify their riboflavin metabolism in an apparent attempt to avoid MAIT cells (35–38), *L. monocytogenes* and a small subset of other pathogens entirely lack the capacity to produce riboflavin and consequently do not synthesize the MAIT cell ligands (39). We propose that the lack of riboflavin biosynthesis provides *L. monocytogenes* with a selective advantage during infection. Interestingly, non-pathogenic members of the *Listeria* genus also require riboflavin, with the exception of *Listeria grayi*, one of the most phylogenetically divergent members of the genus which encodes a riboflavin biosynthesis operon (40). Although the evolutionary history of riboflavin metabolism in the *Listeria* genus remains unclear, we suggest that the absence of the riboflavin pathway represents a pre-adaptive auxotrophic trait that allows *L. monocytogenes* to evade MAIT cells (41).

The results of this study showed that riboflavin-expressing *L. monocytogenes* were recognized by MAIT cells in a MR1-dependent manner both *in vitro* and *in vivo* (**Fig. 2C, 2D, 4E**). At present, it is unclear whether the MR1 ligand is secreted and presented by *L. monocytogenes-ribDEAHT* replicating intracellularly, as is the case in CD8^+^ T cell recognition. It is also possible that the ligand is secreted or released extracellularly since *L. monocytogenes* is known to infect multiple cell types and maintain several extracellular niches during infection (42–44). A recent study showed that intracellular and extracellular bacterial pathogens interact with distinct subsets of immune cells, resulting in different MAIT cell activation and functional phenotypes (45). Identifying the mechanism of *L. monocytogenes*-derived MAIT cell ligand presentation in infected tissues will provide further insight into MAIT cell biology. In a separate study, we showed that *ribDEAHT*-expressing *L. monocytogenes* strains that cannot grow extracellularly still activate MAIT cells, suggesting that intracellular *L. monocytogenes* was primarily responsible for MAIT cell expansion (46).

Previous studies have shown that MAIT cell-derived perforin can restrict bacterial growth *in vitro*, but its role *in vivo* has remained unclear (24, 47). Our data builds on these observations by demonstrating that perforin is required by MAIT cells for both *in vitro* and *in vivo* control of riboflavin-producing *L. monocytogenes* (**Fig. 4)**. In contrast to previous studies that demonstrated that MAIT cell-driven antibacterial response against *Legionella longbeachae* (31) and *F. tularensis* (22) is mediated primarily *via* cytokines (IFN-γ, GM-CSF), our data suggests that MAIT cells rely on perforin to limit riboflavin-producing *L. monocytogenes* growth. However, we cannot exclude that cytokines may also play a role in MAIT cell-dependent antibacterial and antitumor activity in response to riboflavin-producing *L. monocytogenes in vivo*. Indeed, there are MAIT cell subsets that produce different cytokines depending on the immunological contexts resulting in differential MAIT cell functions (48, 49). Furthermore, secreted cytokines can activate other immune cells including NK cells, γδ T cells and other innate-like T cells resulting in distinct immune responses (50, 51). Conversely, MAIT cells may also respond to cytokines present at the infection site (23, 49, 52). Future work will focus on functional characterization of MAIT cell effector functions at different stages of infection in different organs and tumor models.

Infection with WT *L. monocytogenes* synthesizing riboflavin increased the number of MAIT cells in spleens and livers by approximately 5-fold (**Fig. 2**), while Δ*actA* pHyp *ribDEAHT* increased the number of MAIT cells by approximately 20-fold (**Fig. 3B**). Since the Δ*actA* strain is highly attenuated, we used higher infection doses which likely increased infection of antigen presenting cells leading to increased MAIT cell ligand presentation and higher MAIT cell proliferation (19, 36). High doses of Δ*actA* pHyp *ribDEAHT* may also lead to higher levels of IFN-γ, resulting in the production of methylglyoxal, a toxic aldehyde produced by all cells as a byproduct of glycolysis (53, 54). Higher levels of methylglyoxal may in turn increase the production of the MAIT cell ligand (5-OP-RU), which results from a condensation reaction of methylglyoxal and the riboflavin biosynthesis intermediate 5-A-RU (**Fig. 1A**).

MAIT cells accumulate in mucosal tissues of mice infected with *L. longbeachae, Salmonella* Typhimurium, *F. tularensis* and in some models of *Mycobacterium tuberculosis* infection (22, 31, 55, 56). Our results demonstrated that Δ*actA* pHyp *ribDEAHT L. monocytogenes* increased the frequency of MAIT cells in the lungs which was effective at immunization against *F. tularensis* (**Fig. 5C**). We also examined the vaccination efficiency of Δ*actA* pHyp *ribDEAHT L. monocytogenes-*activated MAIT cells in mice lacking perforin and saw no change in protection efficiency (**Fig. S2**). This suggests that the control of *F. tularensis* growth observed in the lungs of WT mice vaccinated with Δ*actA* pHyp *ribDEAHT L. monocytogenes* is not mediated by perforin produced by MAIT cells. We hypothesize MAIT cells in this context might be providing protection via other mechanisms such as cytokine production, which has been observed in both *L. longbeachae* and *F. tularensis* infections (22, 31). Future work will explore the cellular mechanisms of protection mediated by antibacterial MAIT cells and whether immunization with riboflavin-producing *L. monocytogenes* can induce long-term immunity against other riboflavin-producing bacteria.

MAIT cells may display either pro- or anti-cancer functions, but there is increasing interest in using MAIT cells in cancer immunotherapy (57, 58). Here we show that Δ*actA* pHyp *ribDEAHT L. monocytogenes* infection resulted in tumor restriction that was not dependent on CD8^+^ T cells, the primary anti-tumor mediators in Δ*actA L. monocytogenes* cancer models (**Fig. 5E**). MAIT cells can enhance CD8^+^ T cell functions to adenoviral vectors (59), recruit CD8^+^ T cells to the tumor (60), but also inhibit CD8^+^ T cell and NK cell antitumor activity in certain cancer models (61, 62). Although our data show that the frequency of CD8^+^ T cells did not change in the presence of MAIT cells during *L. monocytogenes* infection (**Fig. 3C, Fig. S1**), we cannot rule out the possibility that in the i.v. and i.t. cancer model (33), *Listeria*-triggered MAIT cells may inhibit tumor infiltration of CD8^+^ T cells or suppress their activity. It is also possible that MAIT cells secrete cytokines that activate bystander cells or eliminate immunosuppressive cells in the tumor, such as polymorphonuclear myeloid-derived suppressor cells (33), thereby playing a redundant role with CD8^+^ T cells. This could explain why MAIT cell-mediated tumor-restrictive effects are most evident in the absence of CD8^+^ T cells and why activation of both MAIT cells and CD8^+^ T cells do not synergize and further restrict tumor growth. If the tumor-restricting phenotype in response to Δ*actA* pHyp *ribDEAHT L. monocytogenes* infection is primarily mediated by MAIT cells, it will be important to define the timing of their activation and determine the relative contributions of systemic and tumor-specific MAIT cell populations, as well as other immune cell subsets with known anti-cancer properties across different cancer models (63, 64). Since both MAIT cells and CD8^+^ T cells are prone to undergo exhaustion that abrogates anti-cancer immunity (65–68), sequential activation of CD8^+^ T cells and MAIT cells at different stages of infection may increase the efficacy of *L. monocytogenes* cancer immunotherapies. In addition, some tumors are CD8^+^ T cell-resistant (69) and thus MAIT cell-dependent therapies may fill an important void in the anti-tumor immune arsenal.

The potential use of activated MAIT cells as therapies for cancer and infectious diseases is an exciting area of research where most of the mechanistic insights have come from murine models. However, human MAIT cells can display different effector functions in the periphery compared to MAIT cells from laboratory mice and are more abundant, approximately 10-50-fold higher in peripheral blood and 50-100-fold higher in the liver (70, 71). The high abundance, broad tissue distribution, and capacity of human MAIT cells to rapidly respond to microbial stimuli without the need of patient-specific antigen stimulation, make them attractive candidates for broad-spectrum immunotherapy against infectious diseases and cancer (6, 72). However, MAIT cells in humans have also been linked to inflammatory and autoimmune disorders, although it remains poorly understood whether they are driving tissue pathology, acting as activated bystanders, or attempting to resolve tissue damage (70, 73, 74). Since the role of activated MAIT cells in inflammatory disease appears to be highly context-dependent, the impact of MAIT cell-boosting therapies on patients with inflammatory conditions should be carefully evaluated in future clinical studies. Nevertheless, immunization with well-established safe live bacterial therapies offers a unique approach to treat cancer and could potentially increase the effectiveness of existing cancer therapies, especially those that lose their potency over time by remodelling suppressive tumor microenvironments (63, 75, 76). Further characterization of the interaction between MAIT cells and *L. monocytogenes* expressing riboflavin could offer new and exciting developments in designing treatments against infectious diseases and cancer.

## Materials and Methods

### Bacterial strains and culture

All strains of *L. monocytogenes* used in this study (Table S1) were derived from the WT 10403S strain and were cultured in filter-sterilized brain heart infusion (BHI) media (Becton Dickinson). The strains expressing the *ribDEAHT* operon from a constitutive promoter (pHyp) or its native promoter (pNat) were constructed by amplifying the *ribDEAHT* operon from *B. subtilis* (13, 16) and cloning the constructs into the site-specific pPL2 integrating vector. The plasmids were then introduced into *L. monocytogenes* by conjugation using SM10 *E. coli* (77). Chemically defined synthetic medium was prepared using a previously reported recipe with 1 μM riboflavin (78). Broth growth curves in chemically defined media were generated from *L. monocytogenes* overnight cultures grown at 37 °C with shaking (200 rpm) overnight. The cultures were washed once in 1X sterile PBS, and fresh media was inoculated at an optical density (OD_600_) of 0.03 and OD_600_ was measured every two hours.

### Intracellular macrophage growth curves

Bone marrow from eight-week-old female mice (Jackson laboratory) was used to prepare BMMs. BMMs were cultured in BMM media containing Dulbecco’s modified Eagle’s medium (DMEM), 20% fetal bovine serum (FBS), 10% macrophage colony-stimulating factor, 1% L-glutamine, 1% sodium pyruvate and 14mM 2-mercaptoethanol (Thermo Fisher Scientific). 3×10^6^ BMMs were seeded in 60 mm non-tissue culture-treated dishes (MIDSCI) containing 14 x 12 mm glass coverslips one day prior to infection. *L. monocytogenes* strains were grown overnight without agitation at 30°C. Bacteria were diluted in 1X sterile PBS and infection was performed at a multiplicity of infection (MOI) of 0.25 (79). At 0.5 hours post-infection, BMMs were washed twice with 1X PBS and fresh BMM media was added. 50 μg/mL of gentamicin was added 1 hour post-infection to remove extracellular bacteria. CFUs were determined by collecting coverslips in 5 mL of water and following cell lysis by brief vortexing, the mix was plated on solid agar plates.

### Plaque assays

L2 fibroblasts were cultured in Gibco DMEM (Thermo Fisher Scientific) containing 10% FBS (Avantor-Seradigm), 1mM sodium pyruvate (Corning), and 2mM L-glutamine (Corning). *L. monocytogenes* strains were grown overnight without agitation at 30°C. On the day of the infection the bacteria were diluted in 1X sterile PBS and 1.2×10^6^ L2 fibroblasts per well in 6-well tissue-culture dish were infected at a MOI of approximately 0.1 (80). At 1-hour post-infection, the L2 cells were washed with 1X sterile PBS and overlaid with 1X DMEM containing 0.7% agarose and gentamicin (10 µg/mL), to kill extracellular bacteria, and plates were incubated at 37°C with 5% CO_2_ for 72 hours. L2 cells were then overlaid with a staining mixture containing 1X DMEM, 0.7% agarose, neutral red (Sigma), and gentamicin (10 µg/mL). Following an overnight incubation, the plaque sizes were scanned and analyzed by comparing the diameter of plaques to WT-size plaques using ImageJ (81).

### *L. monocytogenes* virulence experiments

Eight-to-twelve-week-old C57BL/6J mice (The Jackson laboratory), *Prf1^−/−^*KO (The Jackson laboratory), *Mr1^−/−^* KO mice (a kind gift from Dr. Siobhan Cowley at U.S. Food and Drug Administration Division of Bacterial, Parasitic & Allergenic Products), and *Mr1* heterozygous mice (bred inhouse) were infected *via* the tail vein with 200 μL of 1×10^3^ or 1×10^7^ *L. monocytogenes* strains in 1X PBS. Mice were euthanized, and the spleens, livers, and lungs were collected, homogenized in 5 mL, 10 mL and 4 mL of 0.1% IGEPAL (CA-630, Sigma) and plated on BHI medium to determine the number of CFUs per organ.

### *F. tularensis* LVS protection studies

Eight-to-twelve-week-old C57BL/6J mice were infected *via* the tail vein with 1×10^6^ *L. monocytogenes* strains and allowed to recover for 14 days to clear the infection. Mice were then challenged subcutaneously with 1×10^4^ *F. tularensis* LVS. Frozen stocks of LVS, (generously gifted by Dr. Karen Elkins, FDA) were thawed at room temperature and diluted in sterile 1X PBS to the desired concentration prior to infection. At 5 days post-infection with *F. tularensis* LVS, mice were euthanized, and the lungs were harvested and plated on Mueller Hinton plates supplemented with 85 mM NaCl, 1% (w/v) proteose peptone/tryptone, 8% (w/v) bacto-agar, 0.1% glucose, 0.025% ferric pyrophosphate, 2% Isovitalex (#211875, Becton Dickinson, Franklin Lakes, NJ), and 2.5% bovine serum and grown for 3 days at 37°C/5% CO_2_ to determine CFUs of *F. tularensis* LVS.

### Tissue collection and preparation for flow cytometry

For flow cytometry, tissues were collected and processed to obtain single cell suspensions (33, 82). Livers, lungs, or tumors were homogenized to 1 mm^3^ fragments and digested in RPMI 1640 with L-glutamine media supplemented with 25 mM 4-(2-hydroxyethyl)-1-piperazi-neethanesulfonic acid (HEPES), 20 mg/mL DNase I (Roche), and 125 U/mL collagenase D (Roche) using an orbital shaker at 37℃ for 1 hour. Spleens were mechanically disrupted by pressing against a 70-μm nylon mesh filter that was subsequently washed with RPMI 1640 containing L-glutamine. Single cell suspensions from all tissue samples were prepared in ice-cold FACS buffer (PBS with 2% FBS) and subjected to red blood cell lysis using ACK buffer (150 mM NH_4_Cl, 10 mM KHCO_3_, 0.1 mM Na_2_EDTA, pH 7.3) and filtered through a 35 μm nylon mesh filter. Cell surface antigens were stained at room temperature for 30 min using a mixture of fluorophore-conjugated antibodies. Dead cells were stained with Live/Dead Fixable Aqua Dead Cell Stain kit (Molecular Probes). Surface marker staining for murine samples was carried out with anti-mouse CD3 (17A2, BioLegend), anti-mouse CD4 (RM4-5, BioLegend), anti-mouse CD8a (53-6.7, BioLegend), anti-mouse CD45 (30-F11, BioLegend), anti-mouse TCRb (H57-597, eBioscience), anti-mouse TCRgd (GL-3, eBioscience), anti-mouse Thy1.2 (30-H12, BioLegend), and MR1/5-OP-RU Tetramer (NIH Tetramer Core) in 1X PBS. Cells were fixed using the Foxp3/Transcription Factor staining buffer set (eBioscience), prior to intracellular staining. Cells were resuspended in 1X PBS and filtered through a 70-μm nylon mesh filter before data acquisition. Flow cytometry was performed on BD LSR Fortessa X20 (BD Biosciences), CyTEK Aurora (CyTEK Biosciences), or LSRFortessa (BD Biosciences) and datasets were analyzed using FlowJo software (Tree Star). Detailed gating strategy used for the analysis of the specific immune cell populations is presented in **Fig. S4**.

### *In vitro* MAIT cell expansion

Spleens were collected from eight-to-twelve-week-old C57BL/6J and *Prf1*^−/−^ C57BL/6J mice 2-3 weeks following a challenge with 1×10^6^ CFUs Δ*actA* pHyp *ribDEAHT*. Single cell suspensions of spleens were prepared as described above and were incubated with anti-mouse CD4 biotin (RM4-5, BioLegend) and anti-mouse CD8a biotin (53-6.7, BioLegend) to eliminate splenic CD4^+^ and CD8^+^ T cells by negative selection using EasySep magnetic bead kit (STEMCELL Technologies). CD4/8 T cell-free splenocytes were activated with 1 μM 5-A-RU-PABC-Val-Cit-Fmoc (HY-131296, MedChemExpress) in DMEM medium supplemented with 10% FBS, MEM non-essential amino acids (1X) (Gibco), 1 mM sodium pyruvate, 2 mM L-glutamine, 10 mM HEPES, 55 μM β-ME and 200 IU/mL recombinant human IL-2 (TECIMTM, Hoffman-La Roche provided by NCI repository, Frederick National Laboratory for Cancer Research). MAIT cells were cultured for a minimum of 7 days at 37°C with 5% CO_2_ and supplemented with 200 IU/mL IL-2 every 2-3 days. Purity of MAIT cell culture was determined by flow cytometry with Live/Dead Fixable Aqua Dead Cell Stain kit (Molecular Probes), anti-mouse CD3 (17A2, BioLegend), anti-mouse TCRb (H57-597, eBioscience), and MR1/5-OP-RU Tetramer (NIH Tetramer Core) in 1X PBS.

### Co-culture assay of *L. monocytogenes*-infected BMMs with MAIT cells

*In vitro* MAIT cell-dependent cytotoxicity assay was based on the method originally described for CD8^+^ T cell cytotoxicity (30). 3×10^6^ BMMs were seeded in 60 mm non-tissue culture-treated dishes (MIDSCI) containing 14 x 12 mm glass in BMM media overnight one day prior to infection and were used as targeted cells. Prior to co-incubation with BMMs, MAIT cells were expanded *in vitro*, as described above, and were assessed for purity by flow cytometry to determine percentage of MAIT cells of total live cells in heterogenous cultures. BMMs were infected as described above with indicated *L. monocytogenes* strains at a MOI of 0.25. At 0.5 hours post-infection, BMMs were washed 2 times with 1X PBS, supplemented with fresh DMEM, and overlaid with MAIT cells at a 1:1 ratio to infected BMMs at 37°C at 5% CO_2_. At 1 hour post-infection, 50 µg/mL gentamicin (Sigma-Aldrich) was added to the cell culture media to remove extracellular bacteria. Bacterial numbers were determined by collecting coverslips containing infected BMMs and MAIT cells at the indicated time points, followed by cell lysis by brief vortexing in 5 mL of water and plating on BHI agar for CFUs.

### Intravenous and intratumoral (i.v. + i.t.) treatment studies

MC38 cell lines were kindly provided by Dr. Jeff Bluestone’s lab and were maintained in DMEM (GIBCO) supplemented with 10% FBS, 1 mM sodium pyruvate (GIBCO), 10 mM HEPES (GIBCO), and 1 mg/mL penicillin-streptomycin (GIBCO). Tumor cells were grown at 37℃ with 5% CO_2_. For tumor studies, eight-to-twelve-week-old C57BL/6J mice bred in-house were implanted with 5×10J MC38 tumor cells subcutaneously into the shaved right flank. 4 days post-tumor implantation, mice were infected i.v. with 1×10^6^ CFUs of the *L. monocytogenes* strain (either Δ*actA* or Δ*actA* pHyp *ribDEAHT*). At 11 days post-implantation, 5×10J CFUs of the corresponding *L. monocytogenes* strain was injected intratumorally (i.t.) into the palpable tumor. CD8^+^ T cells were depleted by injecting 200 μg of anti-CD8b.2 monoclonal antibody (C2832, Leinco Technologies) intraperitoneally on day 10, 16 and 22 post-tumor implantation. IgG2b isotype control (R1371, Leinco Technologies) was used as a control. Tumor measurements were performed blindly across the entire experiment by a single operator measuring three dimensions of the tumor (width, height and length) every two days starting at 10-12 days post-tumor implantation.

### Animal use and ethics statement

The mice were maintained by University of California, Berkeley Office of Laboratory Animal Care personnel according to institutional guidelines. Animal studies were performed in accordance with the guidance and recommendations of the University of California, Berkeley Office of Laboratory Animal Care, and the Guide for the Care and Use of Laboratory Animals of the National Institutes of Health. The protocols used in this study were reviewed and approved by the Animal Care and Use Committee at the University of California, Berkeley (AUP-2016-05-8811), (AUP-2017-05-9915-2) and protocol no. R353-1113B.

### Statistics and reproducibility

All results were repeated at least twice in independent experiments. All the graphs and statistics were generated using GraphPad Prism (v.10.2.2). For non-normally distributed data (*L. monocytogenes* and *F. tularensis in vivo* infections), the values were log_10_-transformed and one-way ANOVA and Tukey’s multiple comparisons test was performed. The median for all *in vivo* infection data was graphed, and individual CFUs values are represented by dots. For data comparing only two groups, *P* values were calculated with two-tailed *t*-tests using Welch’s correction, as described in the figure legends. For data with more than two comparison groups, ordinary one-way ANOVA with Dunnett’s multiple-comparison test was performed. Two-way ANOVA was performed for data with more than two comparison groups and/or multiple timepoints of measurement. For two-way ANOVA, Šidák’s multiple-comparison correction was used. Tumor volume measurement shows the data of one experiment, as a representative of three independent experiments. Replicate numbers (*n*) are represented in the figures and legends. For all experiments, samples were grouped based on the genotype and treatment group and were not further randomized. Investigators were blinded to experimental groups while performing tumor volume measurements, but not for other experiments. Statistical methods were not used to predetermine sample size.

### Schematics

All schematics were generated in BioRender https://www.biorender.com/.

## Acknowledgments

This work was supported by the National Institutes of Health (grants 1P01 AI063302 [D.A.P.], 1R01 AI027655 [D.A.P.], 5 R01 CA283604 [D.A.P. and M.D.], 1DP2CA247830-01 [M.D.], R01AI113270-01A1 [S.A.S.], R01AI153197-01 [S.A.S.]), the UC CRCC Cancer Research Coordinating Committee (C23CR5612 [M.D.]), Ford Foundation Fellowship and the University of California Dissertation-Year Fellowship (R.R.-L.); the UCSF-UCB SAR Program (M.L.); HHMI Gilliam Fellowship (J.G.C.), CEND fellowship (A.A.-S.). M.D. is a Pew-Stewart Scholar and a St. Baldrick’s Scholar with generous support from Hope with Hazel. We thank the NIH Tetramer Core Facility (NIH Contract 75N93020D00005 and RRID:SCR_026557) for providing mouse MR1/5-OP-RU Tetramer. Portions of this paper were developed from the thesis of R.R.-L.

## Author Contributions

R.R.-L. and D.A.P. conceptualization; R.R.-L., J.G.C., M.L., S.A.S., M.D. and D.A.P. data analysis; R.R.-L., J.G.C., E.T., A.A.-S. and S.E. investigation; R.R.-L., M.L. and D.A.P. writing; S.A.S., M.D. and D.A.P. project supervision; R.R.-L. and D.A.P. funding acquisition.

