## Supplemental figures for "Avoidance of MAIT cells is an essential determinant of *Listeria monocytogenes* pathogenesis"

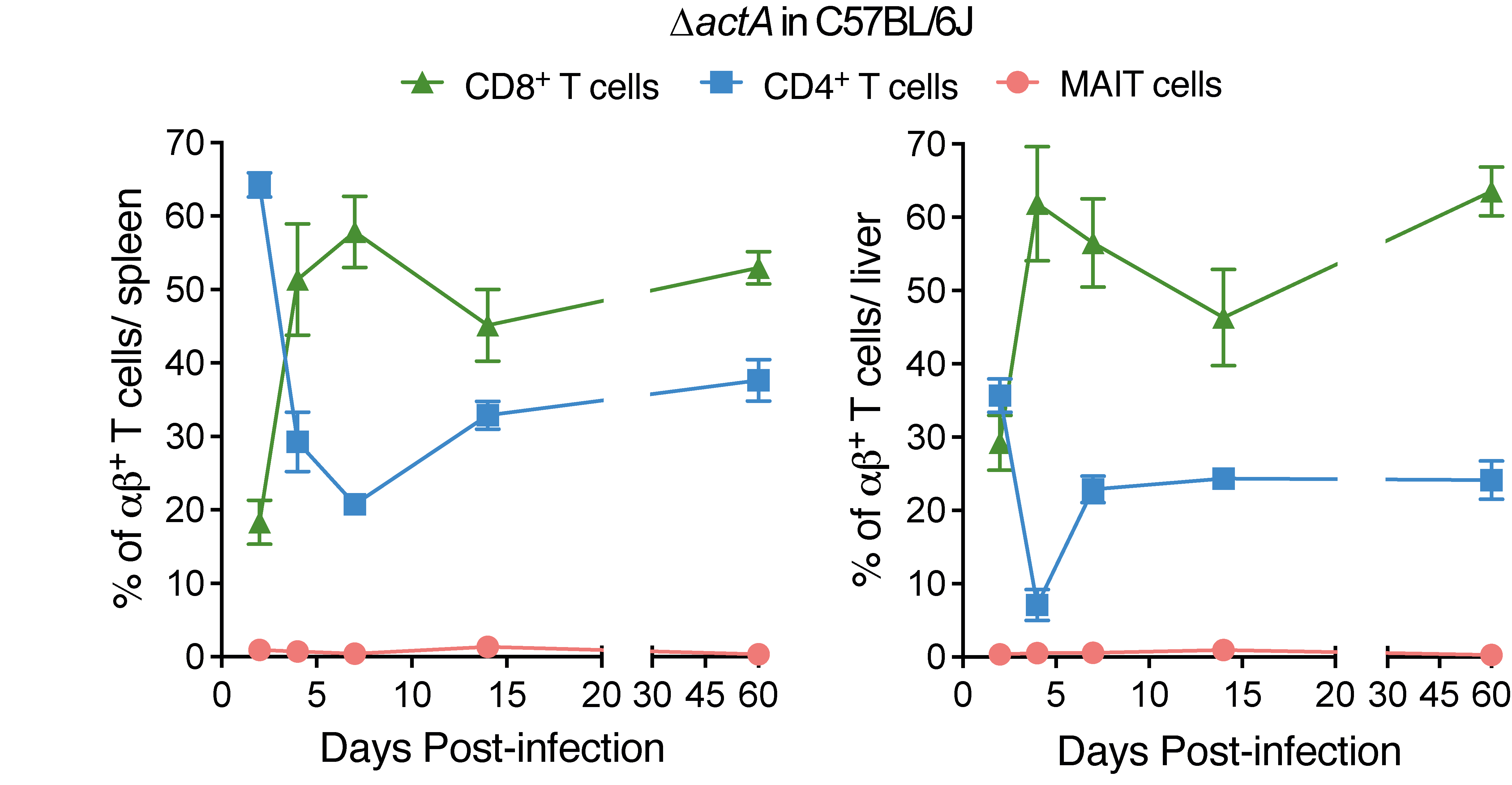

Fig. S1. T cell kinetics during infection with attenuated *L. monocytogenes* Δ*actA* in mice. αβ T cell kinetics in spleens (left) and livers (right) of C57BL/6J mice showing the frequencies of MAIT, CD4^+^, and CD8^+^ T cells at 2, 4, 7, 14, and 60 days post-infection with 1x10^7^ CFUs of Δ*actA.* Data represents two independent experiments with n=10 (day 2), n=4 (day 4), n=7 (day 7), n=6 (day 14) and n=6 (day 60) mice.

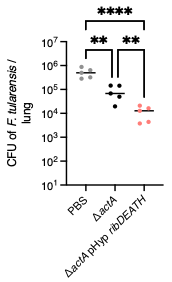

Fig. S2. *Prf1*^-/-^ mice vaccinated with riboflavin-producing *L. monocytogenes* are protected against *F. tularensis* infection. *F. tularensis* LVS lung burdens on day 19 in *Prf1*^-/-^ mice vaccinated with the indicated *L. monocytogenes* strains. PBS (*n*=5), Δ*actA* (*n*=5), and Δ*actA* pHyp *ribDEAHT* (*n*=5). Statistical significance was determined using one-way ANOVA and Tukey’s multiple comparisons test was performed. Means and SEM from one experiment are shown: *n*=5 mice per group. ****P < 0.0001; **P < 0.01.

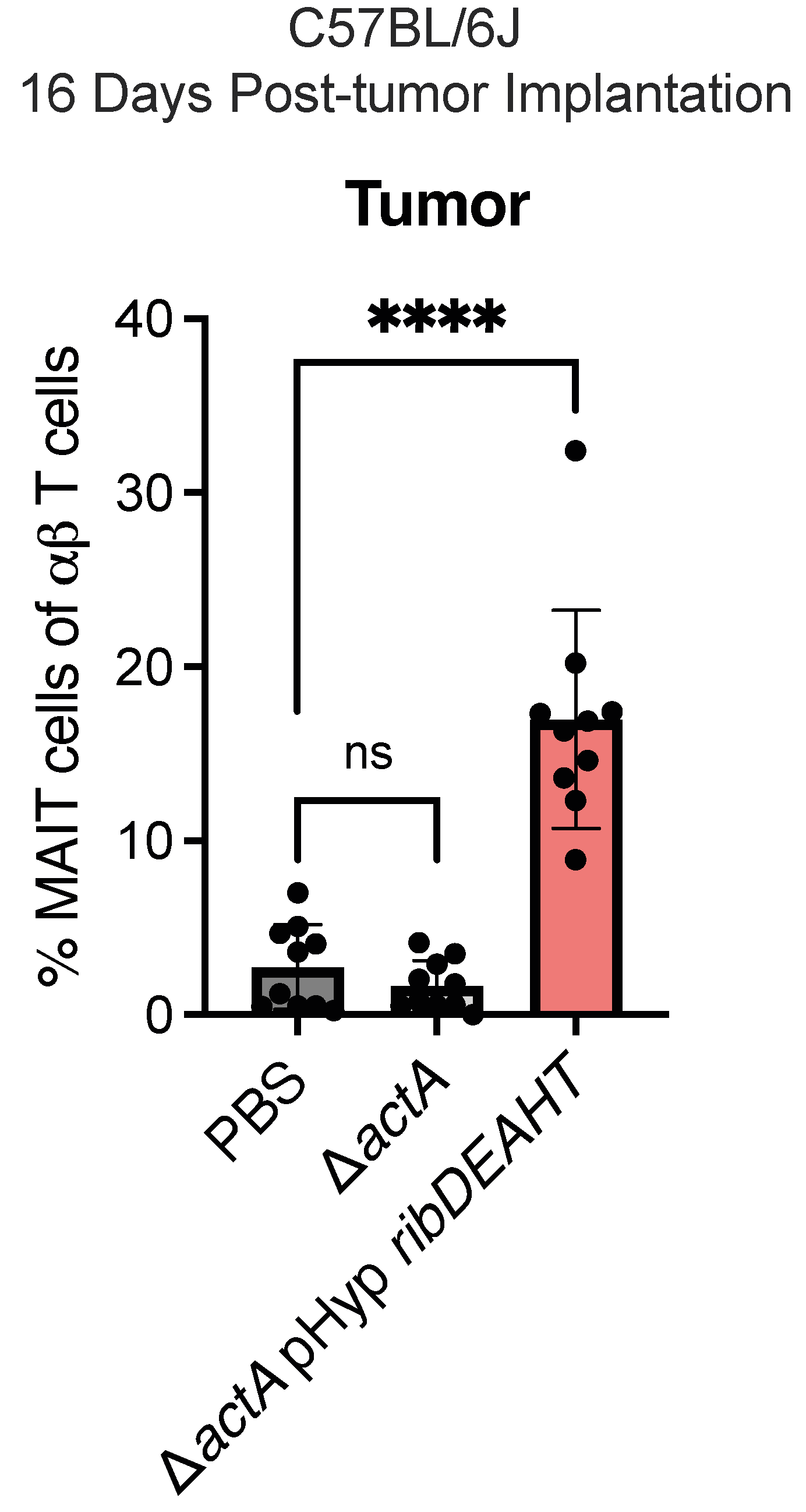

Fig. S3. Frequency of MAIT cells in MC38 tumors of C57BL/6J mice 5 days after i.v. + i.t. treatment with PBS, Δ*actA*, or Δ*actA* pHyp *ribDEAHT*. Means and SEM from two experiments are shown: *n*=10 mice per group. Statistical significance was determined using one-way ANOVA and Dunnett’s post-test with PBS mice as control. ****P < 0.0001; not significant (P > 0.05).

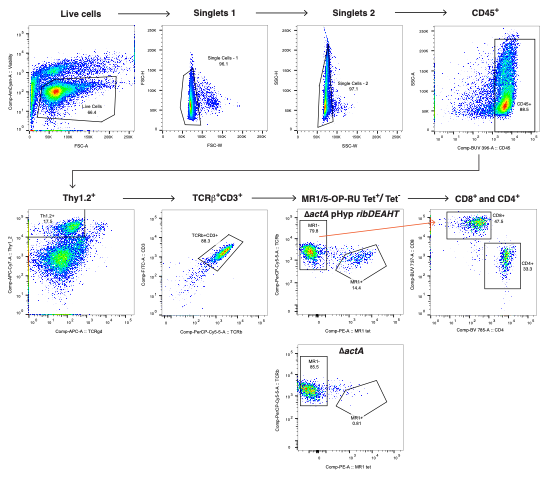

**Fig. S4.** **Gating strategy for flow cytometry analysis**. A representative FACS plot gating strategy used for the analysis of the frequencies of MAIT cells, CD4^+^ and CD8^+^ T cells presented in **Fig 3C**. Same strategy was applied for the analysis of MAIT cell frequency in **Fig. 2B**, **3B**, **4B**, **5A**, and **Fig. S1** and **Fig. S3**.

Tables

Table S1. Bacterial strains used in this study

| **Strain number** | **Background** | **Strain name** | **Reference** |
| --- | --- | --- | --- |
|  | *E. coli* SM10 |  | (1) |
|  | *L. monocytogenes* 10403S | WT | (2) |
| DP-L4029 | *L. monocytogenes* 10403S | Δ*actA* | (3) |
| DP-L7378 | *L. monocytogenes* 10403S | Δ*ribU* pNat *ribDEAHT* | (4) |
| DP-L7490 | *L. monocytogenes* 10403S | pHyp *ribDEAHT* | (5) |
| DP-L7504 | *L. monocytogenes* 10403S | Δ*actA* pHyp *ribDEAHT* | This study |
|  | *F. tularensis* LVS |  | (6) |
